# Friction is preferred over grasp configuration in precision grip grasping

**DOI:** 10.1101/2021.01.13.426550

**Authors:** Lina K. Klein, Guido Maiello, Roland W. Fleming, Dimitris Voudouris

**Author notes:** **Address for Correspondence:** Guido Maiello, Justus-Liebig-Universität Giessen, Abteilung Allgemeine Psychologie, Otto-Behaghel-Strasse 10F, 35394 Giessen, Germany.

## Abstract

How humans visually select where to grasp an object depends on many factors, including grasp stability and preferred grasp configuration. We examined how endpoints are selected when these two factors are brought into conflict: Do people favor stable grasps or do they prefer their natural grasp configurations? Participants reached to grasp one of three cuboids oriented so that its two corners were either aligned with, or rotated away from, each individual’s natural grasp axis (NGA). All objects were made of brass (mass: 420 g) but the surfaces of their sides were manipulated to alter friction: 1) all-brass, 2) two opposing sides covered with wood, while the other two remained of brass, or 3) two opposing sides covered with sandpaper, and the two remaining brass sides smeared with vaseline. Grasps were evaluated as either clockwise (thumb to the left of finger in frontal plane) or counterclockwise of the NGA. Grasp endpoints depended on both object orientation and surface material. For the all-brass object, grasps were bimodally distributed in the NGA-aligned condition but predominantly clockwise in the NGA-unaligned condition. These data reflected participants’ natural grasp configuration independently of surface material. When grasping objects with different surface materials, endpoint selection changed: Participants sacrificed their usual grasp configuration to choose the more stable object sides. A model in which surface material shifts participants’ preferred grip angle proportionally to the perceived friction of the surfaces accounts for our results. Our findings demonstrate that a stable grasp is more important than a biomechanically comfortable grasp configuration.

**NEW & NOTEWORTHY:** When grasping an object, humans can place their fingers at several positions on its surface. The selection of these endpoints depends on many factors, with two of the most important being grasp stability and grasp configuration. We put these two factors in conflict and examine which is considered more important. Our results highlight that humans are not reluctant to adopt unusual grasp configurations in order to satisfy grasp stability.

## INTRODUCTION

When grasping, humans must select appropriate contact points with the object out of a plethora of possible options. This choice is nontrivial and depends on several characteristics of the object, such as its position in relation to the actor (Paulignan et al., 1997; Briere & Proteau, 2011), orientation (Voudouris, Smeets, Brenner, 2013; Paulun et al., 2016), size (Hesse & Franz, 2009; van de Kamp et al., 2009), center of mass (Lukos et al., 2007; Voudouris et al., 2019), surface material (Fikes et al., 1994; Wing & Lederman, 2009), and visibility (Paulun et al., 2016; Maiello, Paulun et al., 2019). We have recently shown computationally that grasp endpoint selection is determined by an intersection of constraints derived from these factors (Klein, Maiello et al., 2020; Maiello, Schepko et al. 2021). Two critical underlying factors for endpoint selection are the prioritization of a stable grasp and the adoption of the natural grasp configuration.

Stable grasps are ensured by applying forces within the cone of friction, so humans bring their digits orthogonally to the object’s surface (Kleinholdermann et al., 2007). When grasping low-friction objects, humans reduce endpoint variability (Paulun et al., 2016) and tailor each digit’s grip forces to the local surface properties (Burstedt et al., 1999), suggesting more careful endpoint selection when anticipating unstable grasps. Unsurprisingly, when grasping elongated objects of combined smooth and rough surfaces, humans choose endpoints on the rough surfaces, presumably to foster grasp stability. Interestingly, though, endpoints are chosen on smooth surfaces if doing so minimizes the torques associated with subsequent object manipulation (Wing & Lederman, 2009; Glowania et al., 2017), suggesting that, although grasp stability is important, other energetic factors are also considered when choosing endpoints.

Grasp control attempts to optimize energy expenditures (Soechting et al., 1995) and minimize travel and spatial error costs (Rosenbaum et al., 2001). A key aspect for selecting the grasp configuration is that extreme joint angles should be avoided (Rosenbaum et al., 2001) because such configurations increase spatial errors (Rosseti et al., 1994). To this end, humans keep their final grasp configurations approximately invariant (Rosenbaum et al., 1992; Grea et al., 2000; Voudouris, Radhakrishan et al., 2013), even when obstacles hinder these configurations (Voudouris et al., 2012a). When grasping cuboid objects that can be grasped with only two configurations, one of which requires the digits to be placed on object positions that are occluded, humans still prioritize their natural grasp configuration by tolerating invisible endpoints (Voudouris et al., 2012b). These examples further highlight the importance of grasp configuration in the selection of endpoints.

Considering the critical role that both grasp stability and final grasp configuration have in grasp endpoint selection, an emerging question relates to the trade-off between these two factors. If grasp stability is prioritized, humans should choose endpoints that provide stable grasps, even when this requires unusual grasp configurations. Alternatively, if final grasp configuration is prioritized, humans should keep their natural grasp posture invariant, even if this would lead to unstable endpoints. To examine this, we asked participants to reach, grasp, and lift cuboid objects of different surface materials. By using cuboids, participants could choose endpoints on only one of the two pairs of opposing surfaces, requiring grasp configurations orthogonal to each other. By manipulating the friction properties of each pair of surfaces, we disentangled the contributions of grasp stability and grasp configuration by examining whether humans prioritize their usual grasp configuration, even if this would sacrifice grasp stability, or whether they prioritize grasp stability by adopting awkward final grasp postures.

## MATERIALS AND METHODS

### Participants

Twenty-one naïve self-reported right-handed participants (mean age: 24.4 years, 16 females) with normal or corrected-to-normal vision participated in our study. This sample size was selected though a-priori power analysis, based on a pilot experiment (N=7), to guarantee that we could detect the smallest effect of interest at the 95% confidence level with 80% power. All procedures were approved by the local ethics board and adhered to the declaration of Helsinki (2013). All participants provided written informed consent prior to the experiment and received monetary compensation for their efforts.

### Apparatus

A schematic depiction of the setup is shown in **Figure 1a**. Participants sat at a table with their right hand at a start position aligned to their shoulder, 9 cm from the table’s edge. Objects were placed in front of the participants at a target position aligned with their midline, 16 cm from the table’s edge. Movements of the participants’ right thumb and index fingers were recorded at 100 Hz with an Optotrak Certus (Northern Digital Inc., Waterloo, ON, Canada) that tracked the position of small infrared markers attached to the respective fingernails (with sub-millimeter accuracy and resolution). A monitor was placed on the table in front of the experimenter, who sat next to the participants. The monitor displayed to the experimenter which condition to set up on each trial. The experiment was programmed in Matlab R2019b using the Motom Toolbox (Derzsi & Volcic, 2018).

**Figure 1:**
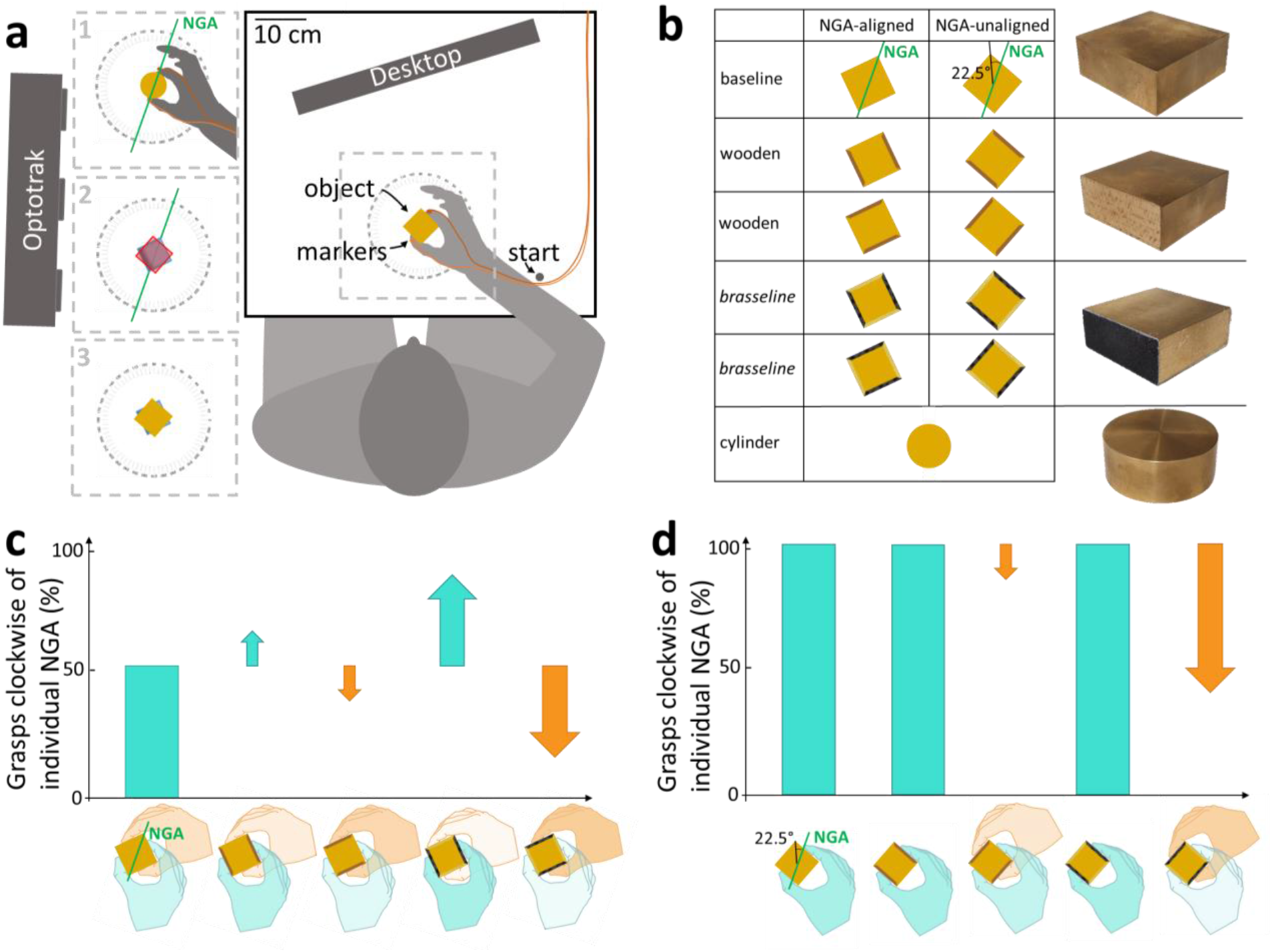
Experimental setup and predictions. **a** Participants reached to grasp and lift an object placed in front of them. After lifting it, they put the object back down, before returning to the start position. **1** They first grasped a cylinder 10 consecutive times to determine each individual’s NGA. **2** The experimenter aligned the object outline template with the NGA. **3** The target cuboid object was aligned with a protractor template according to the condition angle and orientation. **b** Each cuboid was presented at two orientations: with its corners either aligned with the NGA or rotated 22.5° counterclockwise. The wooden and brasseline objects were presented also in two configurations, so that their higher- and lower-friction sides were alternated clockwise and counterclockwise. The cylinder was only used to determine the NGA before the trials involving the cuboids. Object pictures are presented next to the corresponding conditions. **c**,**d** Predictions regarding the percentage of clockwise grasps for different surface material configurations, for **c** NGA-aligned and **d** NGA-unaligned conditions. Arrows visualize a change in behavior compared to the baseline prediction. A thin downwards pointing arrow predicts a small decrease in clockwise grasps, a large arrow a larger decrease. The hands’ degree of translucency represents the amount of predicted clockwise (cyan hand) vs. counterclockwise (orange hand) grasps.

The target objects and the experimental conditions are shown in **Figure 1b**. In the main experiment, the object was one of three possible cuboids (5 cm x 5 cm x 2 cm) that was oriented either with its corners aligned to the participant’s individual NGA or rotated 22.5° counterclockwise. All objects were made of brass (mass: 420 g) but the surface material of the sides was manipulated, so that the sides were either all-brass (*baseline object*), or two of the opposing sides were covered with wood and the other two remained with brass (*wooden object*), or two of the opposing sides were covered with sandpaper and the other two brass sides were made slippery using Vaseline (“*brasseline”* object). Each of the wooden and *brasseline* objects could be placed in two configurations, such that their higher- and lower-friction sides were alternated clockwise and counterclockwise.

### Procedure

Before each trial, participants placed their thumb and index finger at the start position and the experimenter placed the object at the target position at the appropriate orientation and configuration. The experimenter could very precisely position each object at the correct angle by aligning the edges with the corresponding outlines on a protractor template on the table. An auditory cue prompted participants to reach and grasp the object using only their thumb and index finger, and then lift it ∼10 cm high while keeping it level. Participants had to place the object back down at roughly the same position before returning to the start position in anticipation of the next trial. Participants were to execute the task in 3 seconds and could see the object at all times during the experiment. No other instructions were given.

Prior to the main experiment, to measure each individual’s NGA, participants performed 10 grasps to a brass cylinder (diameter 5 cm, height 2 cm, weight 332 g). From these 10 trials, an individual’s NGA was calculated as the median orientation of the grip at the moment of grasp. The experimenter then marked two orientations on the protractor template around the target position so that one corresponded to the calculated NGA (NGA-aligned) and another was rotated 22.5° counterclockwise (NGA-unaligned). Using these outlines, participants performed 6 practice trials drawn from a subset of the experimental conditions, during which they were familiarized with the task and cuboid objects.

The main experiment then started, in which participants grasped only the cuboids. Each of the 10 conditions (**Figure 1b**) was presented 10 times (100 trials per participant), across three object-specific sub-blocks to minimize trial-order effects (Maiello et al, 2018): presentation of baseline, wooden, and the *brasseline* objects were shuffled across participants in a Latin square design. Within each sub-block, object orientation and configuration were presented in pseudorandomized order.

Immediately after the grasping experiment, we asked participants to judge the slipperiness and pleasantness to the touch of each of the four surfaces they could grasp during the experiment. On the monitor participants viewed pictures of the objects with one of the four possible materials facing the participants. Using the mouse in eight separate trials, they first set a slider on a scale from slippery to not slippery and afterwards rated the same four surfaces from pleasant to not pleasant.

### Analyses

#### Endpoints

Endpoints of both fingers with the objects were determined as the coordinates of the markers on the fingernails at the time of contact, as this was determined using the method developed by Schot et al. 2010 and previously described in Paulun et al. (2016). In detail, the average position of the two markers on the fingernails, which represented the position of the hand, had to travel more than half the distance between the start and the target position, and the likelihood of a sample being the moment of contact increased with lower vertical positions and with lower speeds of the hand.

#### Grip Angle

We were interested in which pair of opposing sides was grasped at the moment of object contact in relation to the objects’ surface material. Therefore, for each trial we first computed the final grip angle along the horizontal plane as: *η* = *atan*2(*y*_*index*_ − *y*_*thumb*_, *x*_*index*_ − *x*_*thumb*_). Then, we classified each grip as either clockwise, if the grip angle was *η* < *NGA* (thumb to the left of finger in frontal plane), or counterclockwise, if *η* ≥ *NGA*. Finally, we computed the percentage of clockwise grasps for each participant in each condition.

#### Predictions

Our a-priori, qualitative predictions are illustrated in **Figure 1(c,d)**. For the NGA-aligned orientation there is no obvious preferred grasp configuration for the baseline object (Voudouris et al., 2012b), so participant grasps, at least in the group level, should be split equally between clockwise and counterclockwise. For wooden and *brasseline* objects, we predict more grasps on the wooden and sandpaper sides, respectively, as these surfaces have higher friction and facilitate more stable grasps. For the NGA-unaligned conditions, one pair of sides requires counterclockwise rotations away from the NGA that are twice as large as those required for the clockwise pair of sides (Voudouris et al., 2012b). Therefore, in these conditions we can directly contrast grasp configuration with grasp stability. If the former is more important, participant grasps should be predominantly clockwise, independently of the surface material. If, instead, grasp stability is prioritized, we predict lower proportions of clockwise grasps when the higher friction cube sides are oriented counterclockwise.

#### Statistical analyses: a priori, hypothesis-driven analyses

To assess whether our experimental manipulations shifted participants’ grasps clockwise or counterclockwise, we analyzed the percentage of clockwise grasps using a repeated-measures generalized linear mixed effects model (GLMM) with fixed effects for object orientation, surface configuration, and the interaction between these, plus random subject-level effects. We defined a logit link function and the conditional distribution of the responses as a Binomial distribution. This is conceptually similar to repeated measures analysis of variance, but overcomes ANOVA shortcomings with percentage data (Jaeger, 2008). Comparisons between condition pairs were performed via two-tailed, paired samples t-tests after variance-stabilizing the percentage data via arcsine square root transformation. We report effect size for differences between condition means on variance-stabilized data as *d* = *μ*_*C*1−*C*2_⁄*σ*_*C*1−*C*2_. Statistical significance was set at α < 0.05. All analyses were performed in Matlab version R2019b.

## RESULTS

We investigated grasp endpoint selection when trading-off between grasp configuration and grasp stability. Participants grasped and lifted a cuboid while we varied the surface material of each pair of its sides. This introduced conditions in which higher friction surfaces were orthogonal to the usual grasp configuration, and thereby we could quantify the contribution of each of these factors in endpoint selection.

**Figure 2a** displays each participant’s ten grip orientations and the associated median (the NGA estimate) when grasping the brass cylinder. Across participants, the mean ± standard deviation NGA was 66° ± 9°. **Figure 2b,c** presents an overview of the distributions of the grip angles (relative to each individual’s NGA) for each condition involving the cuboid. For objects aligned with the NGA (**Figure 2b**), baseline grips (top row) were bimodally distributed across participants. This bimodal distribution was somewhat skewed when grasping the low-constraint wooden object (rows 2,3), and became clearly unimodal when grasping the *brasseline* object (rows 4,5). For objects rotated 22.5° away from the NGA (**Figure 2c**), baseline grips were predominantly clockwise, and remained so when the higher friction surfaces were aligned with this natural grasp configuration (rows 2 and 4). Interestingly, when the higher friction surfaces were orthogonal to the baseline grip axis, participants switched their grasp configuration to choose more stable endpoints, subtly for the wooden object (row 3) but massively for the *brasseline* object (row 5).

**Figure 2:**
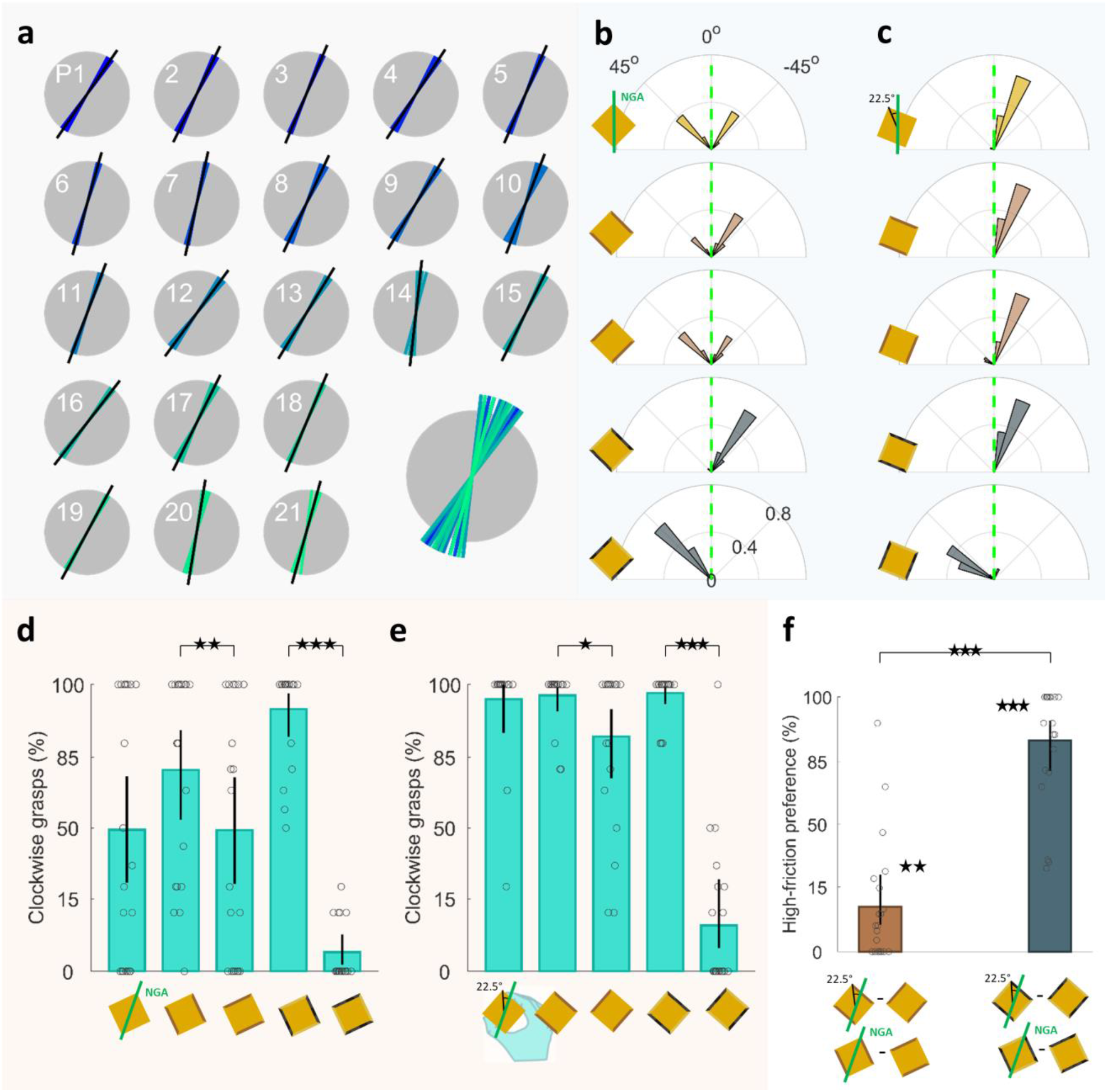
Shifts in grasp orientation following object orientation and surface material. **a** NGA estimation: grip orientation of all ten trials (colored lines) and median grip angles (black lines) for each participant (separate panels) when grasping the brass cylinder. All median NGA estimates are depicted in the lower right panel. **b,c** The proportion of grip angles, relative to each individual’s NGA, for all material configurations. **b** Cube corners aligned with the NGA. **c** Cube corners rotated 22.5° away from the NGA. **d,e** The percentage of clockwise grip angles for each of the five conditions when cuboid corners were **d** aligned with the NGA and **e** when not. **f** Difference in %clockwise grips between surface configurations for the wooden and brasseline objects, collapsed across object orientations. In panels **d,e,f**, circles denote individual participants, bars are means across participants, error bars are 95% bootstrapped confidence intervals. Y-axes are scaled following the arcsine square root transform. *p<.05; **p<.01;***p<.001.

These results were further confirmed by our statistical analyses. GLMM analysis on the percentage of clockwise grasps (**Figure 2d,e**) showed a significant main effect of object orientation (p<.001), as participants grasped the NGA-aligned objects with a bimodal distribution of grip angles, but the NGA-unaligned objects primarily with clockwise grips, in line with previous findings (Voudouris et al., 2012b). The percentage of clockwise grips was further affected by the surface material configuration (p<.001), and this effect was different depending on the object’s orientation (interaction; p<.001). Specifically, for the *brasseline* object, grips were more often clockwise and counterclockwise following the higher friction material in both the NGA-aligned (t(20)=17.9, p<.001, d=3.9) and unaligned orientations (t(20)=13, p<.001, d=2.8). This pattern was observed also for the wooden object, but was weaker (NGA-aligned: t(20)=3.4, p=.0031, d=0.73; NGA-unaligned: t(20)=2.8, p=.011, d=0.61). Note that this interaction arose because in the NGA-aligned orientation, grips shifted both clockwise and counterclockwise from baseline, whereas in the NGA-unaligned orientation they only shifted counterclockwise.

**Figure 2f** further shows the effect of surface material assessed independently of object orientation. Specifically, for each object orientation we calculated the difference in clockwise grasps between the two configurations of each (wooden and *brasseline*) object, and then calculated the average difference across the two object orientations, with greater values indicating stronger preference for higher friction surfaces. We found that grasps were significantly attracted toward the higher friction sides both for the wooden (t(20)=3.4, p=.003, d=0.74) and the *brasseline* objects (t(20)=16.5, p<.001, d=3.6), but that the strength of this attraction was greater in the *brasseline* than the wooden object (t(20)=10.4, p<.001, d=2.3).

Participants performed repeated trials for each condition. We thus wondered whether the observed shifts in grasp orientation were based on visual estimation of object properties or on the memory from repeated experience with the object. To answer this question, we repeated our analyses using only the first trial from each participant in each condition, and found that our findings remained unvaried (correlation between full and reduced dataset: r = 0.99, p<.001).

In short, participants grasped the higher friction surfaces more often than the lower friction surfaces. This was clearly evident when the higher friction surfaces were orthogonal to the baseline grip axis, and particularly apparent in the NGA-unaligned orientation conditions, when grasp stability and final grasp configuration were fully contrasted. Indeed, when grasping the NGA-unaligned *brasseline* object, participants used grasp configurations that were almost never used when grasping the NGA-unaligned baseline object (compare first and last rows of **Figure 2c**). It is possible that participants were content to select these unusual grasp configurations because they could readjust their grip and arm posture when lifting the object off the table. We thus compared grip angles at moment of first contact with grip angles 500 ms after contact, i.e., during object lift. Even when adopting the postures farthest from the NGA (last row of **Figure 2c**), participants readjusted their grip posture on average only by 1 ± 4°, suggesting they maintained postures away from the NGA throughout the grasping action. Therefore, humans prefer endpoints that facilitate stable grasps, even when this requires unusual grasp configurations.

### A simple model: surface friction shifts participants’ preferred grip angle

To gain further insights into the process by which grasp stability and grasp configuration interact when choosing endpoints, we devised a simple model to explain our pattern of results (**Figure 3**). First, we assumed that an individual participant will exhibit a preferred grip axis that follows a normal distribution *N*(*μ*, *σ*), with mean *μ*_*NGA*_ and standard deviation *σ*_*NGA*_. In the equal material conditions (**Figure 3a**), a participant’s grasps will be clockwise or counterclockwise depending on whether this participant’s NGA is clockwise or counterclockwise of the cube’s diagonal, here named *ξ*. Thus, in the NGA-aligned condition (**Figure 3a**, top), where we aligned the cube’s diagonal with each participant’s estimated NGA, approximately 50% of grasps should be oriented clockwise (green shaded region of the distribution) and 50% of grasps should be counterclockwise (orange region). In the NGA-unaligned condition (**Figure 3a**, middle), where the cube diagonal is rotated away from each participant’s measured NGA, most of the NGA distribution should fall clockwise to this diagonal, thus most grasps should be clockwise. The proportion of clockwise grasps *P*_*cw*_ can thus be formalized as the value of the cumulative normal function *Φ*(*x*, *μ*_*NGA*_, *σ*_*NGA*_), evaluated at *x* = *ξ*:

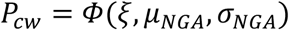

**Figure 3.**
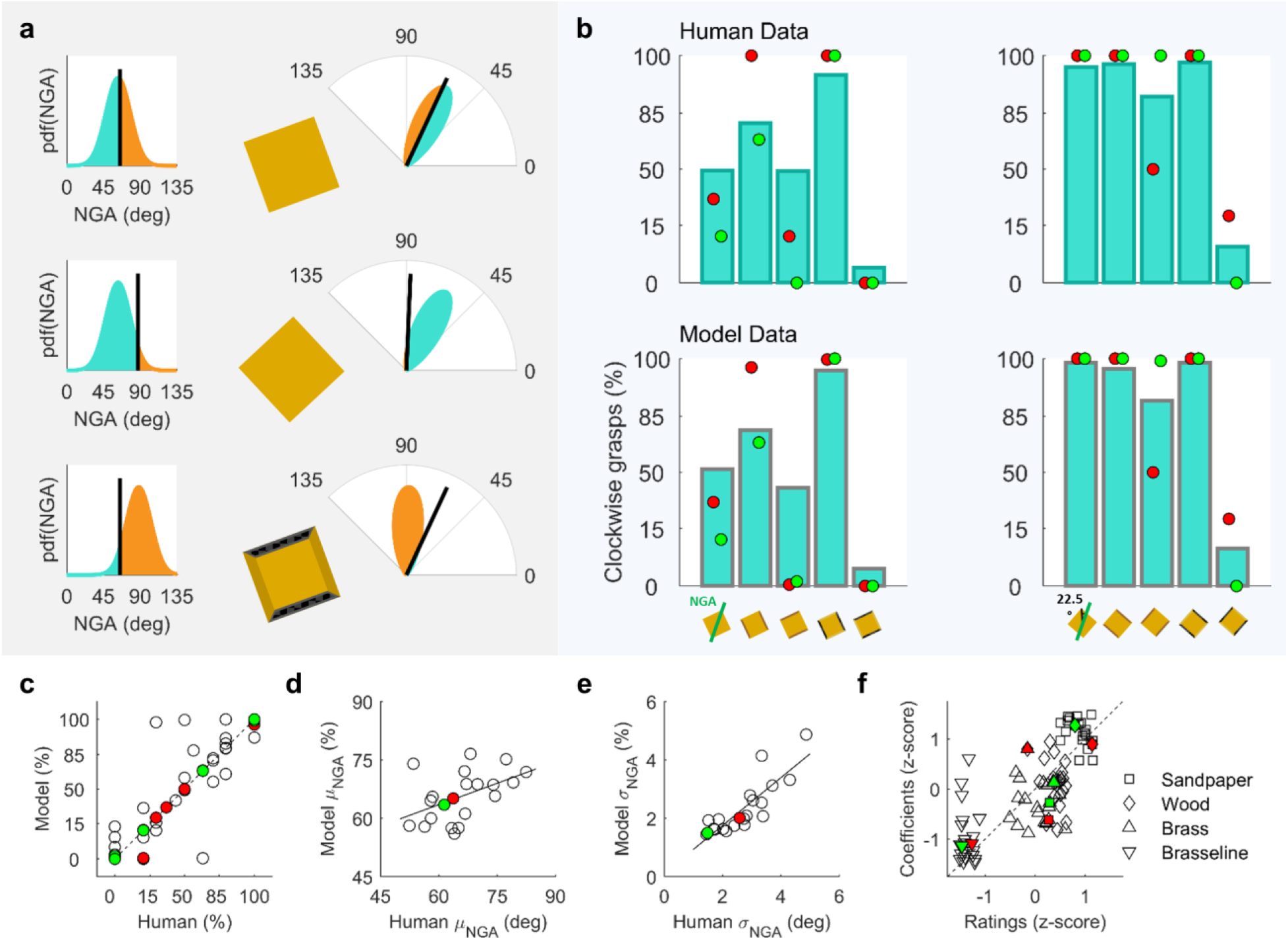
Model results. **a** Model behavior, exemplified at the group level. In the same material conditions (top and middle), the cube diagonals (black lines) in the aligned and unaligned conditions split the NGA distribution into clockwise (green) and counterclockwise (orange) grips by different amounts. For clarity, here we show the NGA distribution in both Cartesian (left) and polar axes (right). In one example condition with different surface materials, the NGA distribution is shifted counterclockwise following the surface with higher friction (sandpaper/black). **b** These shifts very closely capture the patterns of human data, both at the group level, and at the level of individual participants (two example participants are shown as green and red dots). **c** Human vs Fitted model percent clockwise grasps. **d,e** Human vs Fitted model μ_NGA_ and σ_NGA_. **f** Human ratings of perceived surface friction vs Fitted model friction coefficients.

In conditions with different materials at the opposing pairs of surfaces (**Figure 3a**, bottom), we assumed that each individual’s *μ*_*NGA*_ shifts clockwise and counterclockwise by amounts proportional to the perceived friction of the surfaces *v*_*material*_ and to the clockwise or counterclockwise rotations required to grasp these surfaces, *φ*_*θ*,*cw*_ and *φ*_*θ*,*ccw*_, with:

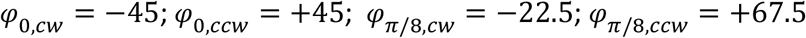

Specifically:

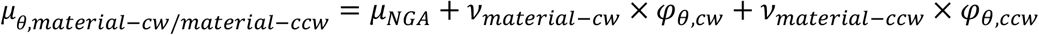

The unknown variables in this framework are thus the positive valued, perceived friction coefficients:

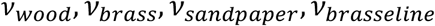

Note that we measured each participant’s NGA prior to our main experiment, and we could thus estimate *μ*_*NGA*_, *σ*_*NGA*_ from these measurements. However, as validation of our model, we also seeded the model with the NGA measurements, but allowed *μ*_*NGA*_, *σ*_*NGA*_ as free parameters. We fit this simple model to each individual participant’s data, and found that the model is able to closely replicate the observed patterns of human data both at the group level (**Figure 3c**) and at the level of individual participants (**Figure 3d**), even after adjusting for the number of predictors in the model (r=0.98, p<.001, r^2^=0.96, r^2^_adjusted_=0.90). Figures 3d and 3e show that the model’s fitted *μ*_*NGA*_ and *σ*_*NGA*_ parameters both significantly correlate with the NGA measurements taken with the brass cylinder object prior to the main experiment (r=0.48, p=.027 and r=0.85, p<.001, respectively). **Figure 3f** further shows that the fitted friction coefficients also significantly correlate with human perceptual ratings of friction (r=0.75, p<.001).

Note that the human perceptual ratings of friction also correlated with the ratings of pleasantness (r=0.80, p<.001), and thus pleasantness ratings also correlated with model friction coefficients (r=0.58, p<.001). However, human perceptual ratings of friction explain 20% more of the variance in the fitted model coefficients.

Given the correlations between model and human NGA parameters, it is also possible to construct a model with only the friction coefficients as free parameters, fixing *μ*_*NGA*_ and *σ*_*NGA*_ to the experimentally measured values for each participant. This reduced model is also able to replicate the observed patterns of human data (r=0.84, p<.001, r^2^=0.71, r^2^_adjusted_=0.51), and the fitted friction coefficients again significantly correlate with human perceptual ratings of friction (r=0.65, p<.001) better than with pleasantness ratings (r=0.45, p<.001). This simple model is thus able to directly relate human perception of surface friction to the selected hand posture for grasping.

## DISCUSSION

We examined whether the selection of grasp endpoints depends primarily on the grasp stability or on the adoption of the usual grasp configuration at the moment of the grasp. To this end, we first measured individual NGAs and then oriented cuboid objects so that their corners were either aligned with the individual NGA or rotated 22.5° counterclockwise from the NGA. By having participants grasp cuboids, we implicitly asked them to grasp the cuboid object with one of two possible grasp configurations. By placing these cuboids at two different orientations, we created conditions in which grasps could be either bimodally distributed or systematically directed to one pair of the object’s sides. By further manipulating the materials of the object’s surfaces, we introduced conditions in which the axis connecting the higher friction surfaces was orthogonal to the grasp axis required for adopting the usual final grasp configuration, eventually allowing us to test which of the two factors is more important for contact point selection. Our results are clear: Humans choose endpoints that promote stable grasps, even if this requires adopting unusual grasp configurations.

The object’s orientation influenced the selection of endpoints as expected (Voudouris et al., 2012b). When grasping the NGA-aligned all-brass object, grasp orientations at the population level were bimodally distributed, reflecting that objects could indeed be grasped from both pairs of sides without adopting awkward grasp configurations. Interestingly however, single participant grasps were less bimodally-distributed than at the group level, even when participants should not have had a clear preference in grip orientation, perhaps reflecting the fact that grasp planning, such as grip forces and digit placement, is sensitive to sensorimotor memories obtained in previous trials (Lukos et al., 2013; Witney et al., 2000). When grasping the NGA-unaligned all-brass object, the grasp distribution was clearly unimodal, suggesting that participants systematically chose grasp configurations within the midrange of their joints and avoided extreme joint angles at the moment of the grasp (Rosenbaum et al., 2001), likely to avoid pronounced endpoint errors (Rosseti et al., 1994).

Our main interest, though, was whether participants would sacrifice their usual grasp configuration to choose stable endpoints or whether they would tolerate endpoints on the lower friction surfaces to maintain their usual grasp configuration. We show that participants were content to adopt unusual grasp configurations that foster grasp stability. This is reflected in the systematic switches of grasps between the two different configurations of each (wooden and *brasseline*) object, and is highlighted in the clear change of behavior when grasping the different configurations of the *brasseline* object: Participants tailored their grasp configurations to ensure that their digits landed almost always on the sandpaper rather than on the vaseline-covered surface (see **Figure 2d,e**). This behavior was also evident for the wooden object, but less pronounced. A possible reason for this difference might be that participants avoided the vaseline-covered surface for other reasons than slipperiness per se, for instance to avoid having vaseline stuck on their digits or due to the unpleasantness of that material. We believe that this is unlikely, as our modelling demonstrates that participant grasps are more directly related to perceived surface friction, rather than perceived pleasantness to the touch. Rather, the difference between the wooden and the *brasseline* objects should be attributed to the lower relative costs of grasping the two surfaces of the wooden object (brass over wooden) compared to the greater costs of grasping the two surfaces of the *brasseline* object (vaseline-covered brass over sandpaper). Of course, if grasping the higher friction surfaces required particularly extreme grasp configurations (e.g., at the very limits of what is biomechanically possible), participants may have favored the lower friction of the two alternative grasps, as long as they could produce sufficient forces to overcome the lack of friction. However, we find that within the range of conditions tested, participants spontaneously adopted grasp configurations that they otherwise would almost never produce to avoid the difficulties associated with grasping a slippery surface.

Our model linking perceptual ratings of friction to final grip orientation hints at how a simple neural circuit could implement these changes in grip selection in the brain, for example within the network formed between the Ventral Premotor Cortex (Area F5), Dorsal Premotor Cortex (Area F2), and the Anterior Intraparietal Sulcus (AIP). Areas F5 and F2 encode grip-wrist configuration and orientation (Raos et al, 2004; Raos et al, 2006). Both regions exhibit strong connections with AIP (Murata et al, 2000), which in turn plays a key role in linking the ventral visual stream (where visual material properties are encoded) to the hand motor system (Borra et al, 2008). Therefore, through area AIP, estimates of surface friction coming from ventral visual areas could bias our preferred grip orientation encoded in areas F5 and F2.

Choosing grasp endpoints requires the consideration of several factors. Two main factors are grasp stability and the final grasp configuration (Klein, Maiello et al., 2020). Grasp stability is important when controlling grasping and choosing endpoints (Paulun et al., 2016; Smeets & Brenner, 1999). Interestingly, humans have been found to sacrifice grasp stability in order to adopt grasp configurations that minimize other energy-related costs, such as torques during object manipulation (Glowania et al., 2017). Yet the magnitude of grip force that is required to overcome surface friction also determines energy expenditure and thus places additional constraints on grasp point selection, suggesting a crucial role of surface material properties in grasping. By directly juxtaposing the contributions of grasp configuration and stability, we demonstrate that participants systematically chose endpoints that promote stable grasps, even when these endpoints required grasp configurations that would otherwise be avoided. We conclude that humans strive for stable grasp endpoints at the expense of their final grasping posture.

## ACKNOWLEDGMENTS

This research was supported by the DFG (IRTG-1901: “The Brain in Action” and SFB-TRR-135: “Cardinal Mechanisms of Perception”), and an ERC Consolidator Award (ERC-2015-CoG-682859: “SHAPE”). Guido Maiello was supported by a Marie-Skłodowska-Curie Actions Individual Fellowship (H2020-MSCA-IF-2017: “VisualGrasping” Project ID: 793660).

## DATA AVAILABILITY

Data and analysis scripts will be made available from the Zenodo database (doi: 10.5281/zenodo.xxxxxxx upon publication).

## Notes

### Competing Interest Statement

The authors have declared no competing interest.

